# iSHARC: Integrating scMultiome data for heterogeneity and regulatory analysis in cancer

**DOI:** 10.1101/2025.04.28.651068

**Authors:** Yong Zeng, Shalini Bahl, Xi Xu, Xinpei Ci, Tina Keshavarzian, Lin Yang, Ho Seok Lee, Pathum Kossinna, Federico Gaiti, Gregory W. Schwartz, Housheng Hansen He, Mathieu Lupien

## Abstract

**Summary:** The 10x Genomics single cell Multiome (scMultiome) assay enables the simultaneous profiling of chromatin accessibility and gene expression from the same nucleus, and has increasingly been utilized in revealing cellular heterogeneity and gene regulation in cancers. However, a dedicated bioinformatics pipeline specifically designed for this type of data is still lacking. Here we present iSHARC, a streamlined pipeline for quality control, modality integration, clustering, cell type annotation, and regulatory mechanism analysis of individual scMultiome data, as well as for integrating multiple samples. The main advantages of iSHARC are: 1) easy implementation, execution and extension through a modular Snakemake workflow management system; 2) flexible analysis and parameters customization via a single configuration file; and 3) comprehensive accessibility by providing different access points to results and detailed summary reports from a single run.

**Availability and implementation:** This pipeline is an open-source software under the MIT license and it is freely available at https://github.com/yzeng-lol/iSHARC.

**Supplementary information:** Supplementary data are appended.

## 1 Introduction

In contrast to standalone single-nucleus ATAC-seq (snATAC-seq) and single-cell/nucleus RNA-seq (sc/snRNA-seq), simultaneous profiling of epigenomic and transcriptomic landscapes from the same cell nucleus enables the identification of novel cell types or states characterized by the features from both modalities, as well as provides the most direct linkage between DNA regulatory elements and their target genes (Ma *et al*., 2020). In late 2020, 10x Genomics introduced the Chromium scMultiome ATAC + Gene Expression (RNA) sequencing protocol, allowing for simultaneous measurements of accessible chromatin status and gene expression from the same nuclei (Shi *et al*., 2021). This technology has increasingly been applied in revealing cellular complexity and gene-regulatory dynamics in human cancers (Foster *et al*., 2022; Liu *et al*., 2023; Derrien *et al*., 2023; Terekhanova *et al*., 2023). However, a dedicated bioinformatic pipeline specifically tailored for scMultiome data is still lacking in the field. Here we present iSHARC, a Seurat-centered (Stuart *et al*., 2019; Hao *et al*., 2021) pipeline that provides a streamlined workflow for quality control, modality integration, clustering, cell type annotation, and regulatory mechanism analysis of individual samples. It also supports the integration of multiple samples by first integrating each individual modality using three different strategies, followed by modality integration. iSHARC was developed with Snakemake (version 7.25.0) (Mölder *et al*., 2021) following a modular design, ensuring automatic installation, seamless execution, and easy incorporation of new analysis modules. It offers flexibility by allowing users to customize analysis inclusion and parameter settings via a single configuration file. Additionally, a single iSHARC run offers access to Seurat objects compiled from initial to extended analysis results, ensuring adaptability to evolving research needs and objectives. It also generates summary HTML reports that provide a comprehensive overview about the data quality and primary findings, facilitating further exploration and investigation.

## 2 Pipeline description

iSHARC consists of two primary modules for analyzing individual samples and integrating multiple samples. For each individual sample, the pipeline starts with 10x Genomic scMultiome ATAC + RNA FASTQ data, followed by data preprocessing and QC, integrating modalities and clustering, annotating integrated cell clusters, and revealing regulatory mechanisms (**Fig. 1** and **Supplementary Fig. S1**). For multiple samples, the workflow first employs three strategies–simple merging, harmonization and anchoring–for horizontally integrating ATAC and RNA data across samples separately, followed by vertical integration of both modalities (**Fig. 1** and **Supplementary Fig. S1**). The pipeline parses a single input configuration file in YAML format to apply user-customized analysis and parameter settings (**Supplementary Table 1)**. Detailed instructions and a template for the configuration file are available in the GitHub repository. All outputs from iSHARC are organized in corresponding folders (**Supplementary Table 2 and 3**). Notably, iSHARC generates a summarized HTML report for QC and primary results for both individual samples and integrated samples (**Fig. 1**), providing an overview of the data and facilitating further exploration and investigation.

**Figure 1.**
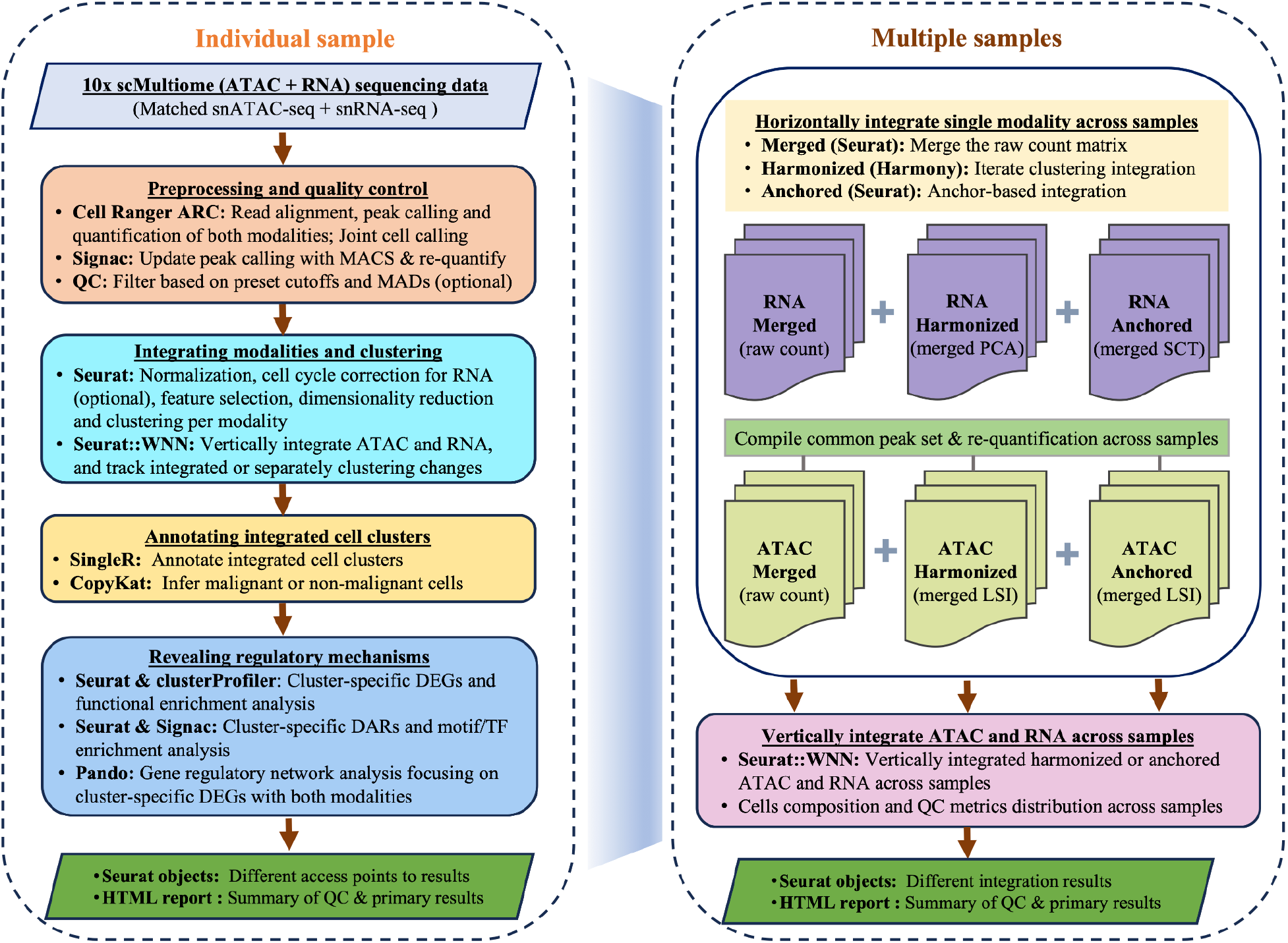
Flowchart of the iSHARC pipeline. Abbreviation: QC, quality control; MAD, median absolute deviation; DEGs, differentially expressed genes; DAR, differentially accessible regions; TF, transcription factor. PCA, principal component analysis; SCT, sctransform normalization in Seurat; LSI, latent semantic indexing.

### 2.1 Preprocessing and QC

First, iSHRAC employs Cell Ranger ARC (version 2.0.2, 10x Genomics) for read alignment, ATAC peak calling, quantification of both ATAC and RNA data, and filtering of background noise through joint cell calling (**Fig. 1**). This step is skipped if iSHARC starts with preprocessed data from Cell Ranger ARC. Next, iSHARC refines ATAC peak calling using MACS2 (version 2.2.9.1) through the CallPeaks function of the Signac (version 1.11.0) (Stuart *et al*., 2021), followed by re-quantification and QC assessments for ATAC data. Additionally, iSHARC offers an optional second round QC, which can further remove low quality cells by taking preset thresholds (Terekhanova *et al*., 2023) and the median absolute deviation (MAD) of selected QC metrics into account. The QC metrics examined for each individual sample include the number of transcripts per cell (nCount_RNA) and percent of reads originating from the mitochondria (pct_MT) for RNA data, as well as the number of unique nuclear fragments (nCount_ATAC), TSS enrichment score, and nucleosome signal for ATAC data (**Supplementary Fig. S2A, B**). However, since these QC metrics can vary substantially between sample types, particularly cancer samples (Heumos *et al*., 2023), users are encouraged to set more customized thresholds as needed. To support this, iSHARC also calculates selected percentiles for these QC metrics to assist users in determining their thresholds (**Supplementary Fig. S2C**).

### 2.2 Integrating modalities and clustering

Next, iSHARC utilizes Seurat (version 4.3.0.1) for normalization, feature selection and dimension reduction for both ATAC and RNA data **(Fig. 1**). During the normalization process for RNA data, iSHARC provides an option for cell cycle assessment and correction (**Supplementary Fig. S2D**). However, it is recommended to deactivate this option if the samples are undergoing differentiation processes as suggested by the Seurat documentation. Subsequently, iSHARC applies the weighted-nearest neighbor (WNN) method implemented in Seurat v4 (Hao *et al*., 2021) to vertically integrate ATAC and RNA modalities (Lee *et al*., 2023). Uniform manifold approximation and projection (UMAP) embedding and clustering are then performed for integrated modalities, as well as each modality independently (**Supplementary Fig. S3A**). In addition to evaluating the modality weight for each integrated cluster (**Supplementary Fig. S3B**), iSHARC tracks the changes in clustering groups for each cell using integrated modalities or separately via an interactive Sankey plot (**Supplementary Fig. S3C**), enabling users to identify instances where RNA-based clusters split into subclusters after integrating the ATAC profiles, or vice versa.

### 2.3 Annotating integrated cell clusters

Although standalone ATAC and RNA profiles generally identify similar cell types, joint profiling can reveal novel cell types or status that exhibit discordance between the two modalities (Ma *et al*., 2020; Lee *et al*., 2023). To capitalize on this, iSHARC focuses on the cell clusters derived from the integrated ATAC and RNA embedding (**Fig. 1)**. The integrated clusters are automatically annotated using SingleR (version 1.4.1), which leverages the compiled reference transcriptomes for pure population of stroma and immune cells generated by Blueprint and ENCODE (**Supplementary Fig. S4A**) (Aran *et al*., 2019). In addition, to distinguish cancer cells from non-malignant cell types, iSHARC employs, CopyKAT (version 1.1.0), which uses integrative Bayesian approaches to calculate DNA copy number events from RNA data without needing a reference set of normal cells (Gao *et al*., 2021). In this analysis, cells predicted as diploid and aneuploid are considered normal and tumor cells, respectively (**Supplementary Fig. S4B**).

### 2.4 Revealing regulatory mechanisms

Additionally, iSHARC identifies both cluster-specific differentially expressed genes (DEGs) and differentially accessible regions (DARs) by comparing each cluster with all other clusters using Seurat’s FindMakers function (**Fig. 1**). For each group of DEGs, the functional enrichment analyses, including Gene Ontology (GO) and Kyoto Encyclopedia of Genes and Genomes (KEGG) pathway analysis, are performed to provide further insights into their biological significance (**Supplementary Fig. S5A, B**). In addition, the top 5 upregulated DEGs per cluster are visualized in a heatmap (**Supplementary Fig. S5C**), and the associated ATAC peaks linked to these DEGs are also exported (**Supplementary Table 2**). For cluster-specific DARs, motif enrichment analysis is conducted on the top-ranked DARs with an adjusted p value smaller than 0.005 using Signac’s FindMotifs function. The JASPAR2020 database (version 0.99.10) of transcription factor (TF) binding profiles (Fornes *et al*., 2020) is used, and the top 5 enriched motifs and related transcription factors per cluster can also be visualized in a heatmap (**Supplementary Fig. S5D**).

The paired scMutiome data also enables more accurate inference of cellular gene regulatory networks (GRN), unveiling the interplay between chromatin, TFs, and genes specific to particular cell types or states (Badia-I-Mompel *et al*., 2023). iSHRAC incorporates Pando (version 1.0.4), which infers GRNs by modeling the expression of each gene based on both TF expression and the accessibility of their putative binding site by using paired ATAC and RNA data (Fleck *et al*., 2023). Specifically, iSHARC constructs a network between TFs and combined integrated cluster-specific genes as targets (**Supplementary Fig. S6A**), as well as cluster-specific GRNs between TFs and individual groups of cluster-specific genes (**Supplementary Fig. S6B**). iSHARC customized GRN graphs allow users to easily distinguish TFs and their targets and the direction of regulation, whether activation or inhibition (**Supplementary Fig. S6**), facilitating further experimental assessment and validation.

### 2.5 Integrating multiple samples

scMutliome cancer studies often involve samples from multiple patients or those collected under varying conditions, such as different tissue sources, treatments or capture times. Therefore, iSHARC provides a snapshot of sample differences by examining the distribution of cell numbers, cell phase and type compositions (**Supplementary Fig. S7A, B**), and QC metrics across samples (**Supplementary Fig. S7C**). Furthermore, these scMultiome datasets are often generated at different experimental times or by different personnel, making batch effect evaluation and correction essential when integrating data from multiple samples (Tran *et al*., 2020). To integrate scMultiome data across samples, iSHARC applies a two-step process: horizontal integration of each individual modality, followed by vertical integration of both modalities (Argelaguet *et al*., 2021). For horizontal integration, iSHARC initially merges all data to assess batch effects, then applies both Harmony (version 1.2.3) (Korsunsky *et al*., 2019) and Seurat RPCA (Stuart *et al*., 2019; Hao *et al*., 2021) for batch correct correction and integration across samples (**Supplementary Fig. S8A, B**). These tools are benchmarked and recommended as primary methods for most scenarios (Tran *et al*., 2020). The WNN method is then applied for the vertical integration of both modalities, which have been processed by merging, Harmony and Suerat, respectively (**Supplementary Fig. S8C, D**). This step is followed by examining the consistency of annotated cell types within clustering groups based on the integration of multiple samples (**Supplementary Fig. S9)**. Together, these processes enable the users to assess and select the most appropriate integration strategy. Lastly, regulatory mechanisms analysis can also be easily performed using the same functions designed for individual samples, facilitated by the consistent data structure post-integration.

## 3 Implementation

Detailed instructions on how to install and run iSHARC, along with templates for the configuration file and submission script for parallelization on a high-performance computing system, are available at (https://github.com/yzeng-lol/iSHARC). To demonstrate the pipeline’s functionality, we utilized a publicly available lymphoma scMultiome dataset from 10x Genomics (https://www.10xgenomics.com/datasets/fresh-frozen-lymph-node-with-b-cell-lymphoma-14-k-sorted-nuclei-1-standard-2-0-0) for individual sample analysis (**Supplementary Fig. 2-6**). For multiple sample integration, we used in-house scMultiome data from three metastatic prostate cancer rapid autopsy samples upon request (**Supplementary Fig. 7-9**). The corresponding HTML reports for summarizing their QC and primary results are available at (https://github.com/yzeng-lol/iSHARC/tree/main/test).

## Acknowledgement

We acknowledge the Princess Margaret Genomics and Bioinformatics group for providing the infrastructure required to conduct analyses included in this work.

## Supplementary Data

### Supplementary Figure

**Supplementary Figure 1.**
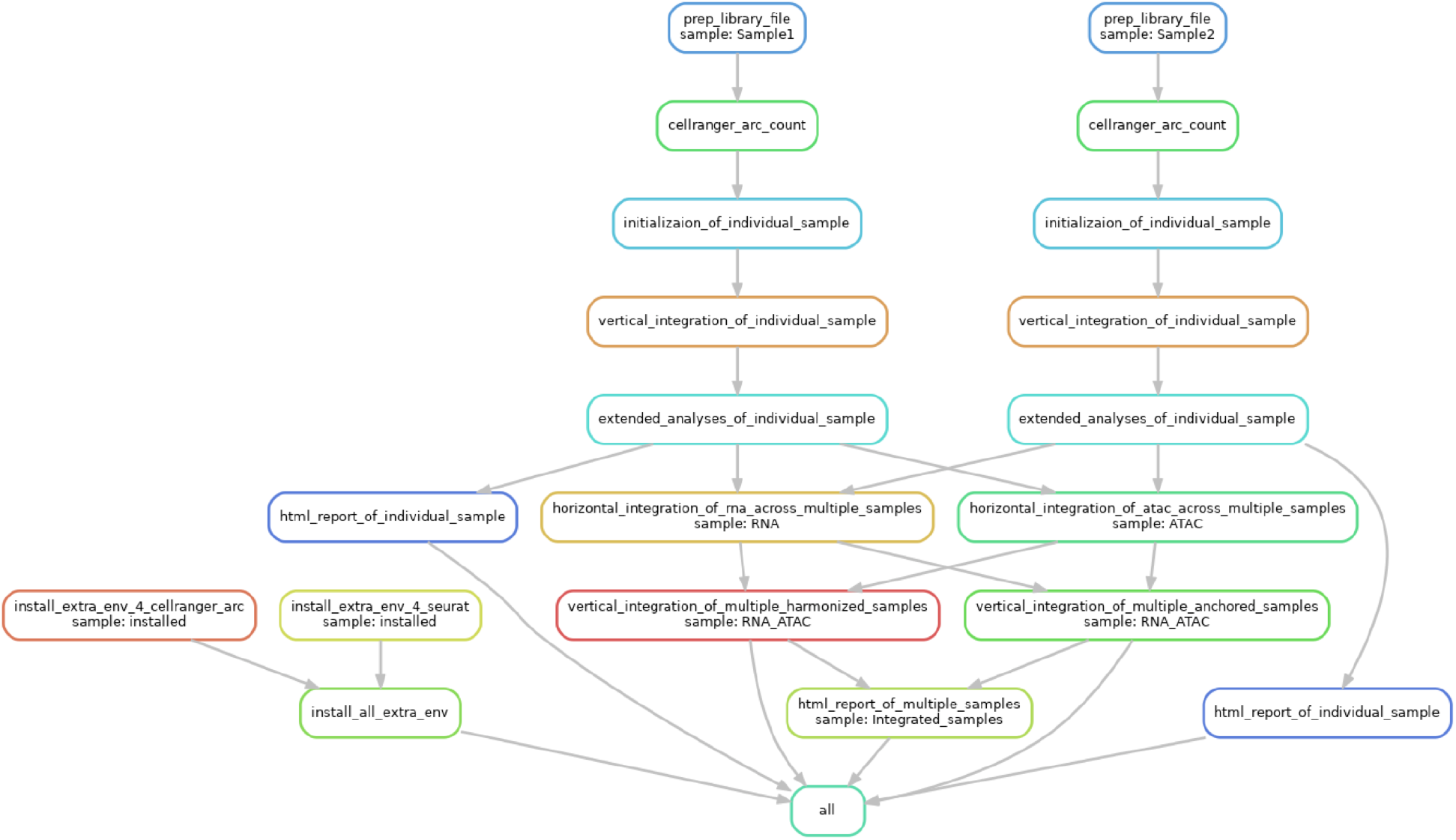
The Snakemake directed acyclic graph (DAG) illustrating the dependencies between rules in the iSHARC workflow.

**Supplementary Figure 2.**
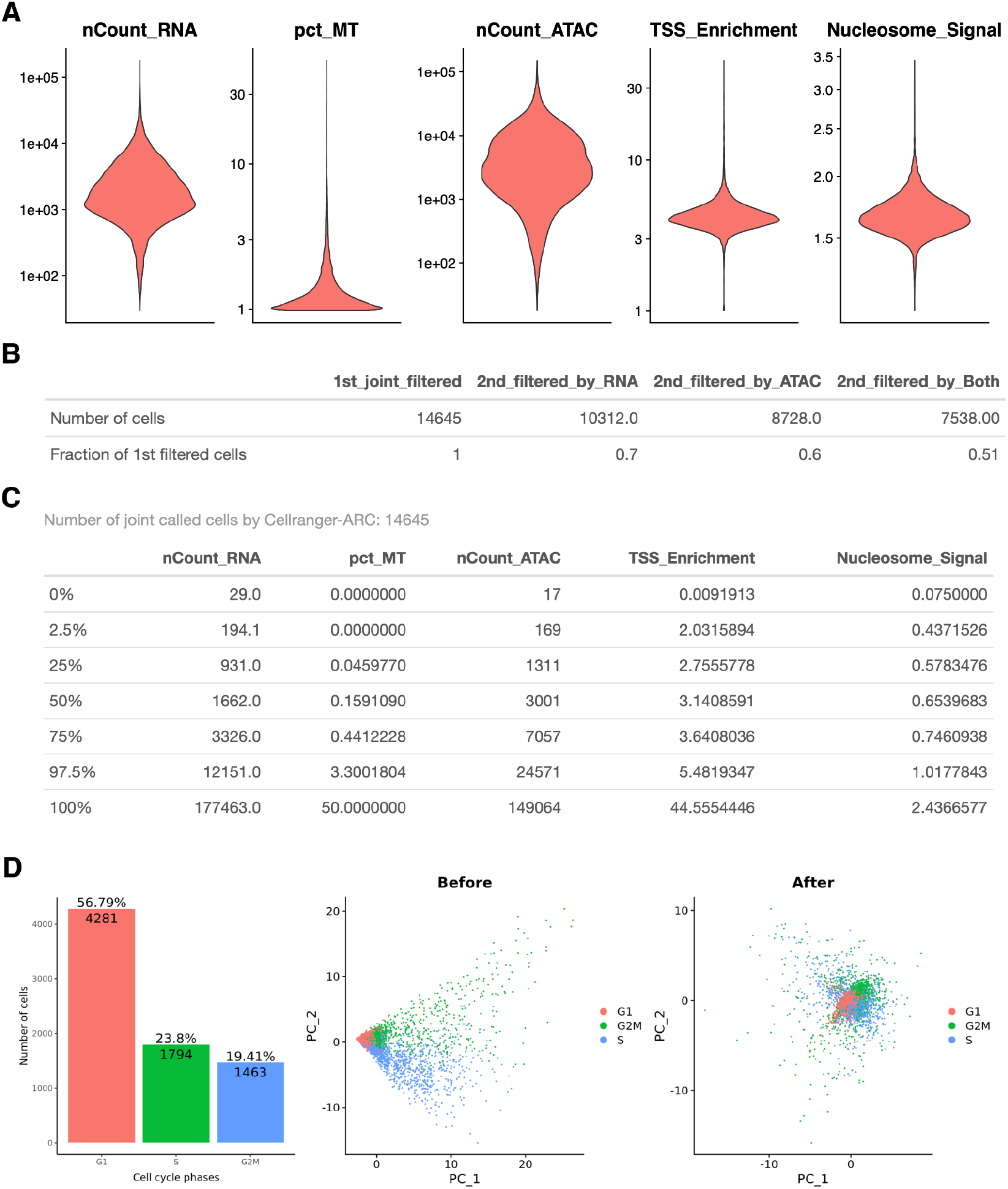
**A)** Violin plots displaying the distribution of selected QC metrics. **B)** The assessment of second round QC filtering by RNA, ATAC alone and together. **C)** Percentile thresholds for selected QC metrics. **D)** The number of cells in the G1, S and G2M phases of the cell cycle (left), and PCA plots showing the distribution of cells before and after cell cycle correction, where only cell cycle-related genes were included in the PCA analysis.

**Supplementary Figure 3.**
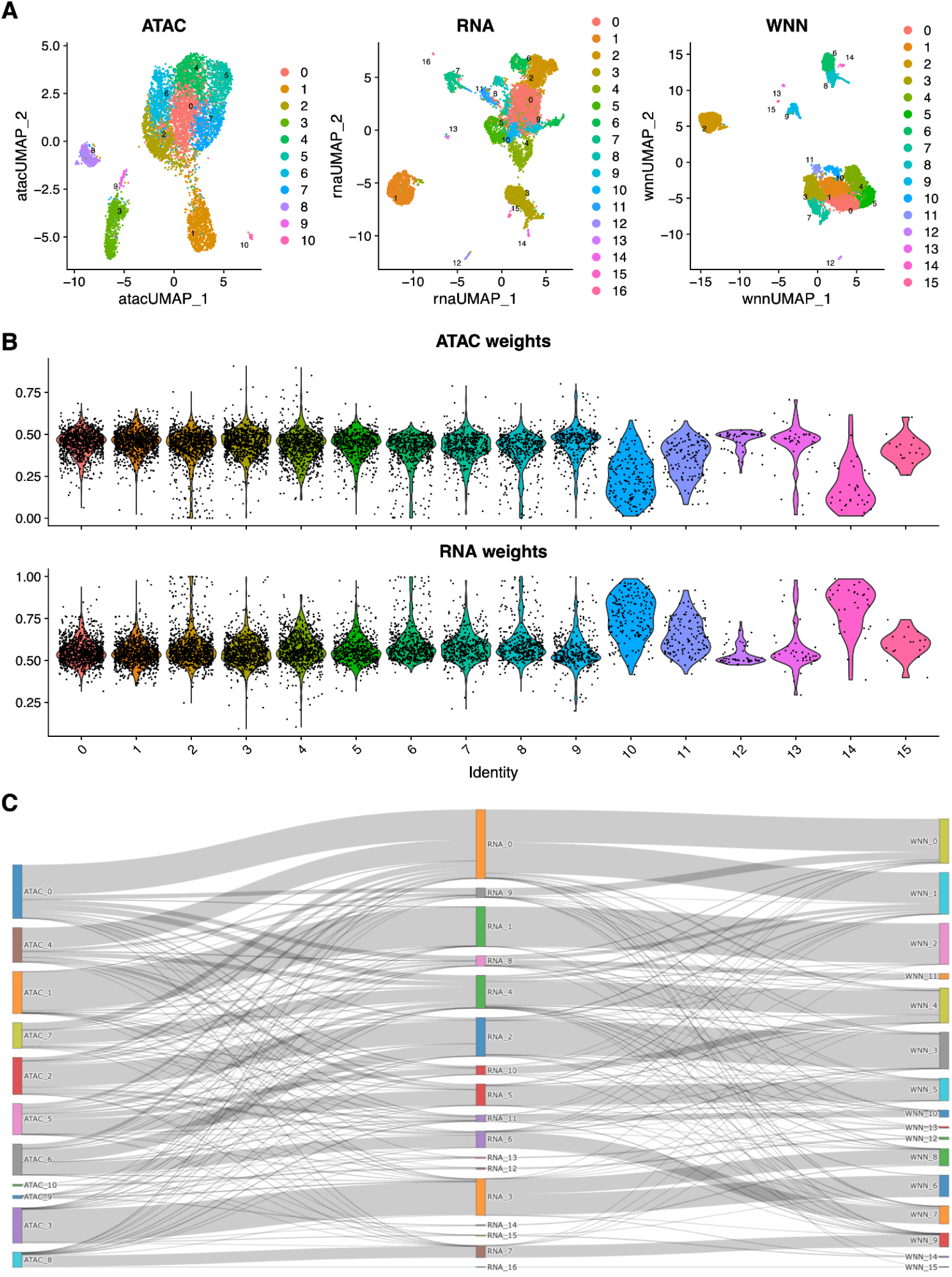
**A)** UMAP plots showing clustering results for ATAC, RNA, and WNN integrated modalities. **B)** Modality weights reflect the relative contribution of ATAC and RNA for each WNN integrated cluster. **C)** Sankey plot illustrating the transitions between clusters identified using ATAC, RNA, and WNN integrated data, with more information interactively readable in the iSHARC HTML report for individual samples.

**Supplementary Figure 4.**
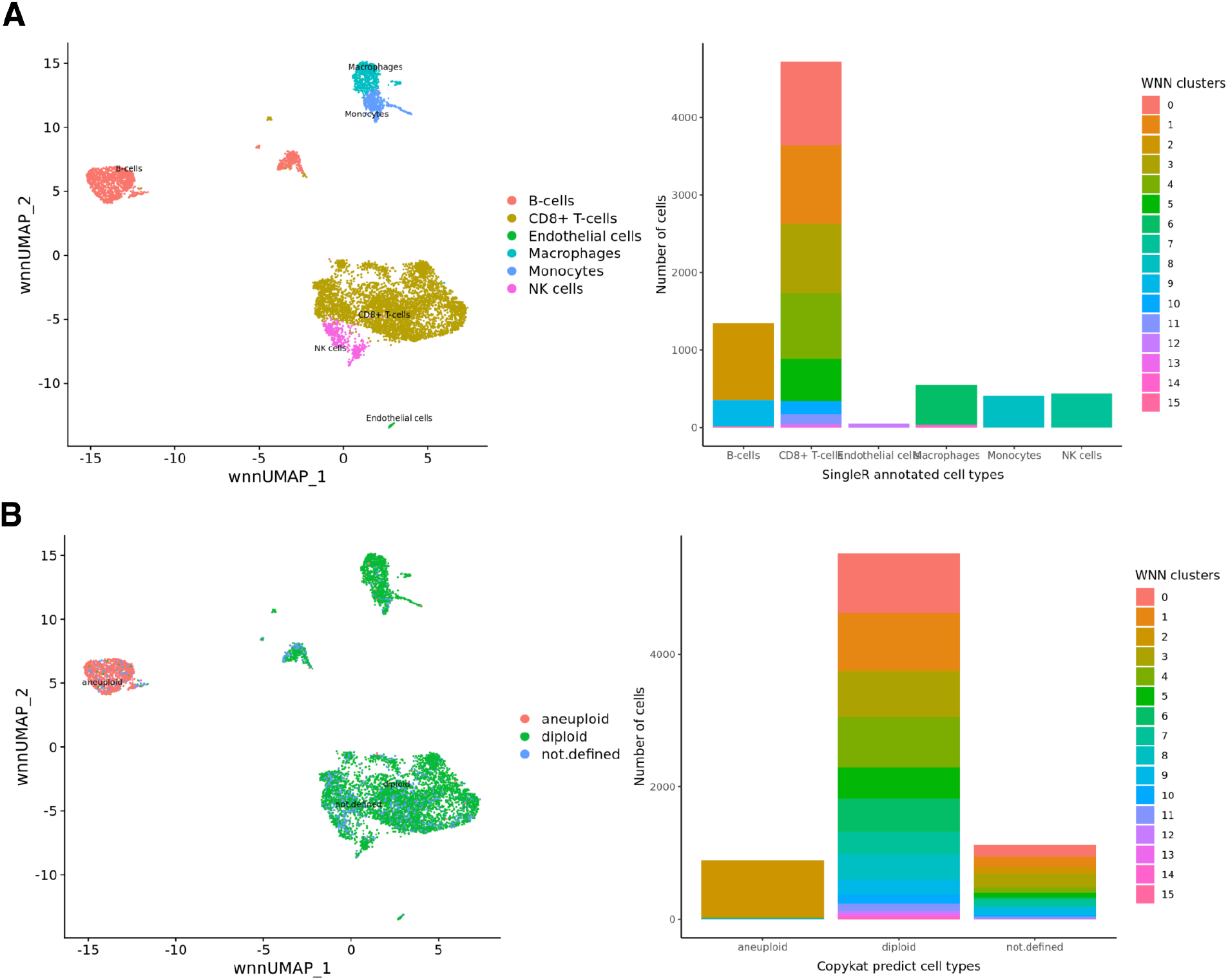
**A)** UMAP plot labeled with SingleR-based cell type annotations (left), alongside the distribution of cells for each annotated cell type across integrated clusters (right). **B)** UMAP plot labeled with cell type annotations inferred by copyKat (left), alongside the distribution of cells for each annotated cell type across integrated clusters (right).

**Supplementary Figure 5.**
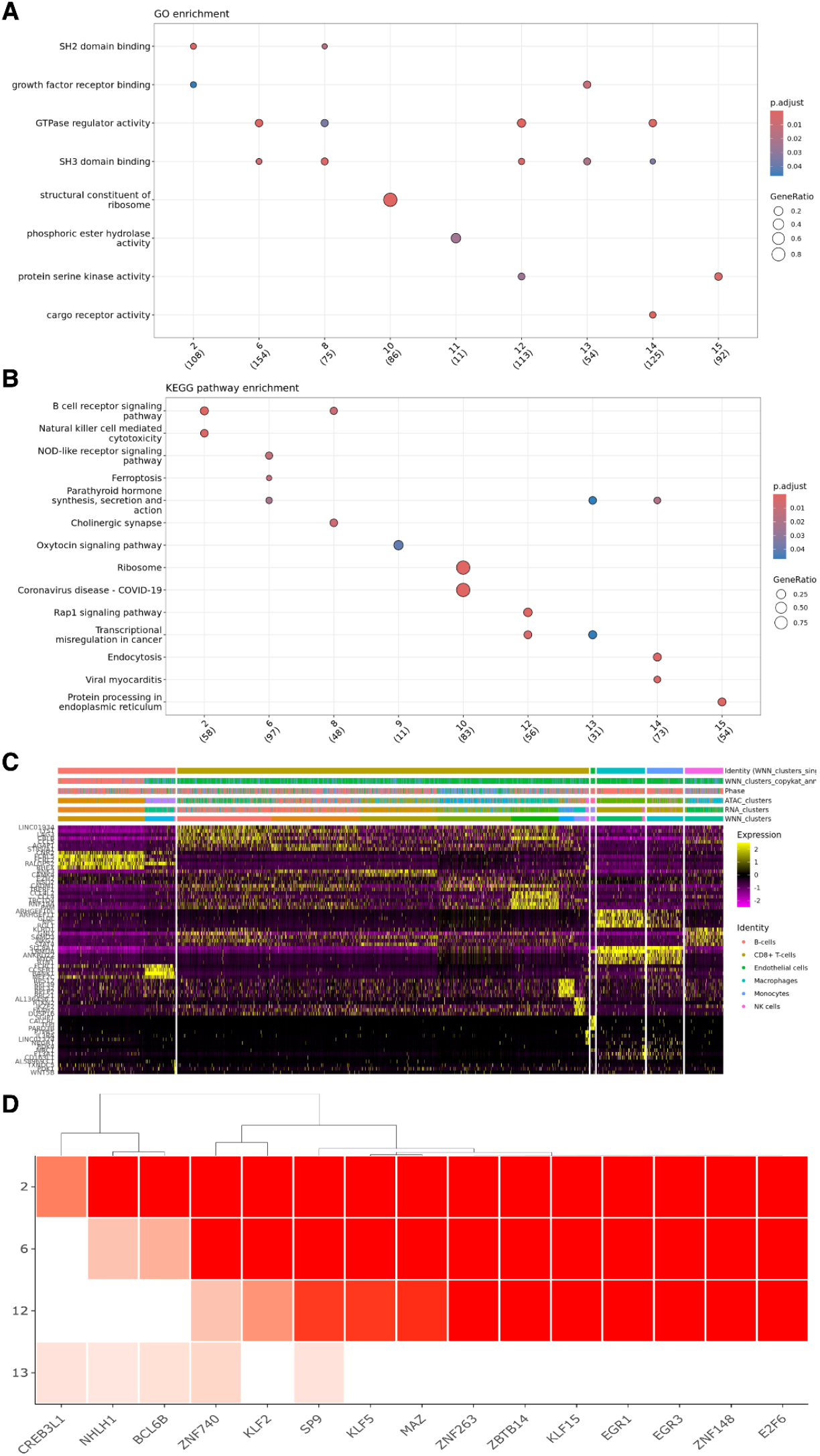
Enriched GO terms **(A)** and KEGG pathways **(B)** for cluster-specific DEGs. **C)** Heatmap showing the top 5 upregulated cluster-specific DEGs, with top color bars indicating different clustering groups and cell type annotations. **D)** Heatmap displaying the top enriched TFs associated with the top cluster-specific DARs, with p-values interactively readable in the iSHARC HTML report for individual samples.

**Supplementary Figure 6.**
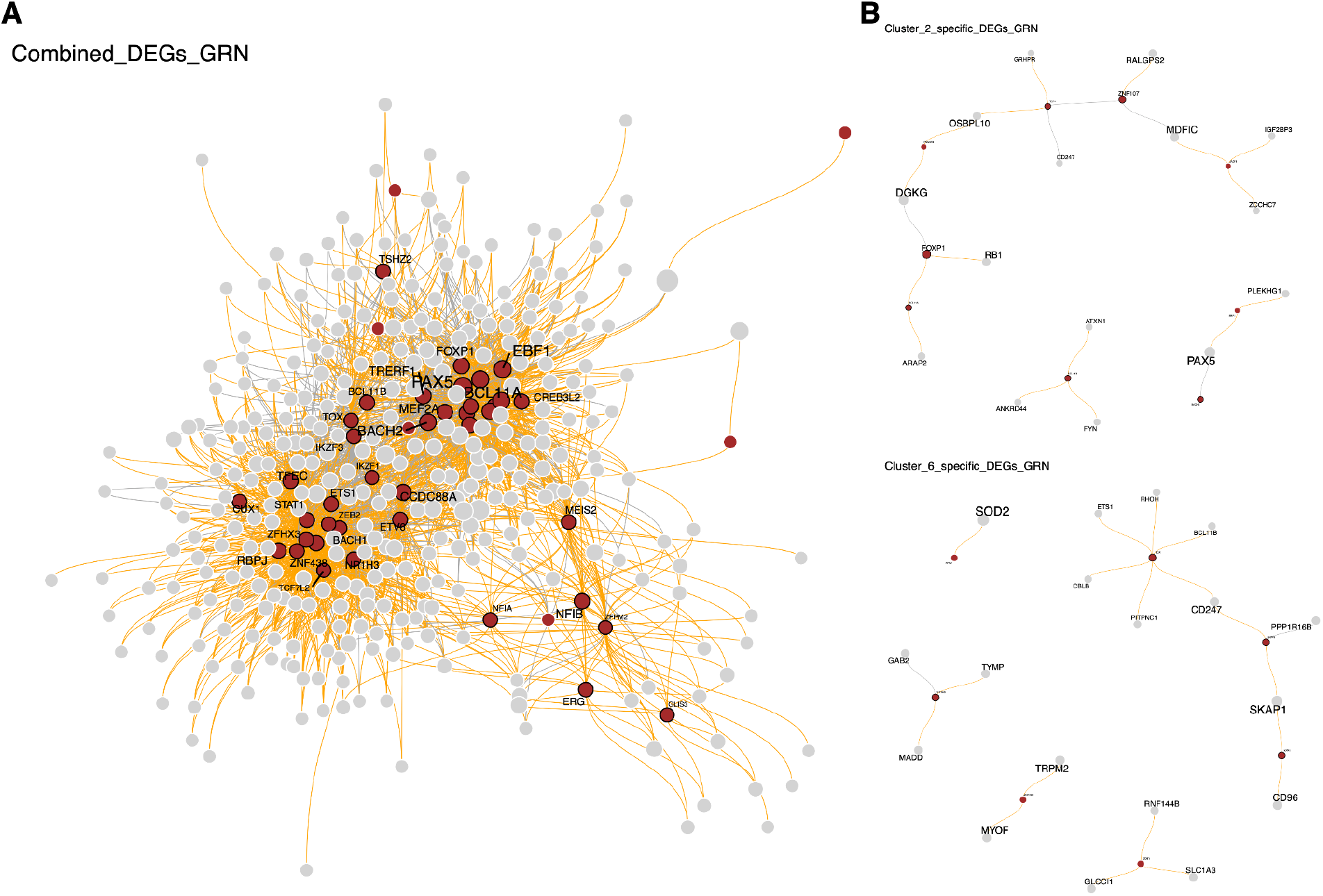
**A)** Gene regulatory network based on all combined integrated cluster-specific DEGs. **B)** Cluster-specific gene regulatory networks based on corresponding cluster-specific DEGs.

**Supplementary Figure 7.**
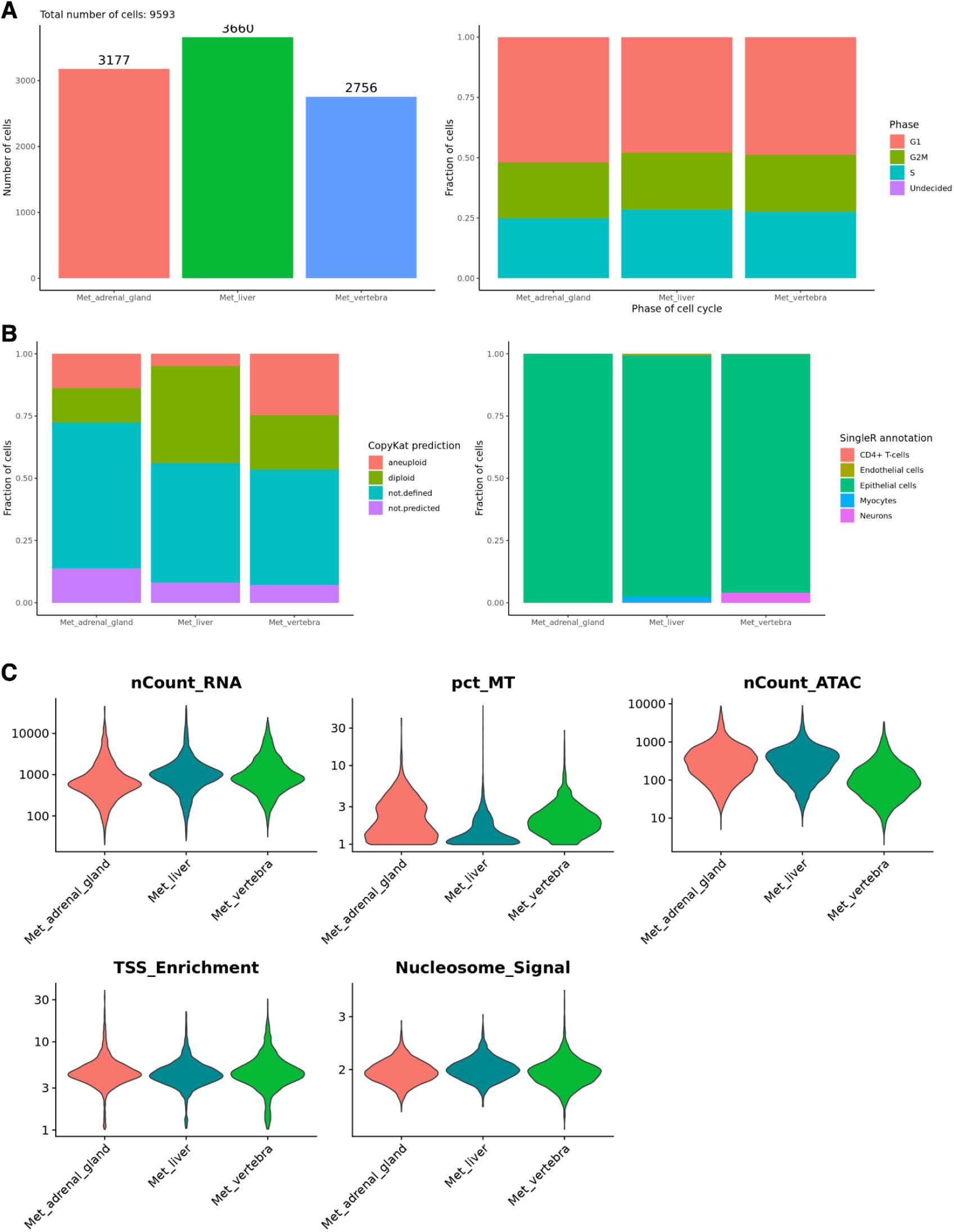
**A)** Number of cells across samples (left), along with the fraction of cells in different cell cycle phases (right). **B)** Fraction of different annotated cell types across samples. **C)** Violin plots showing the distribution of selected QC metrics across multiple samples.

**Supplementary Figure 8.**
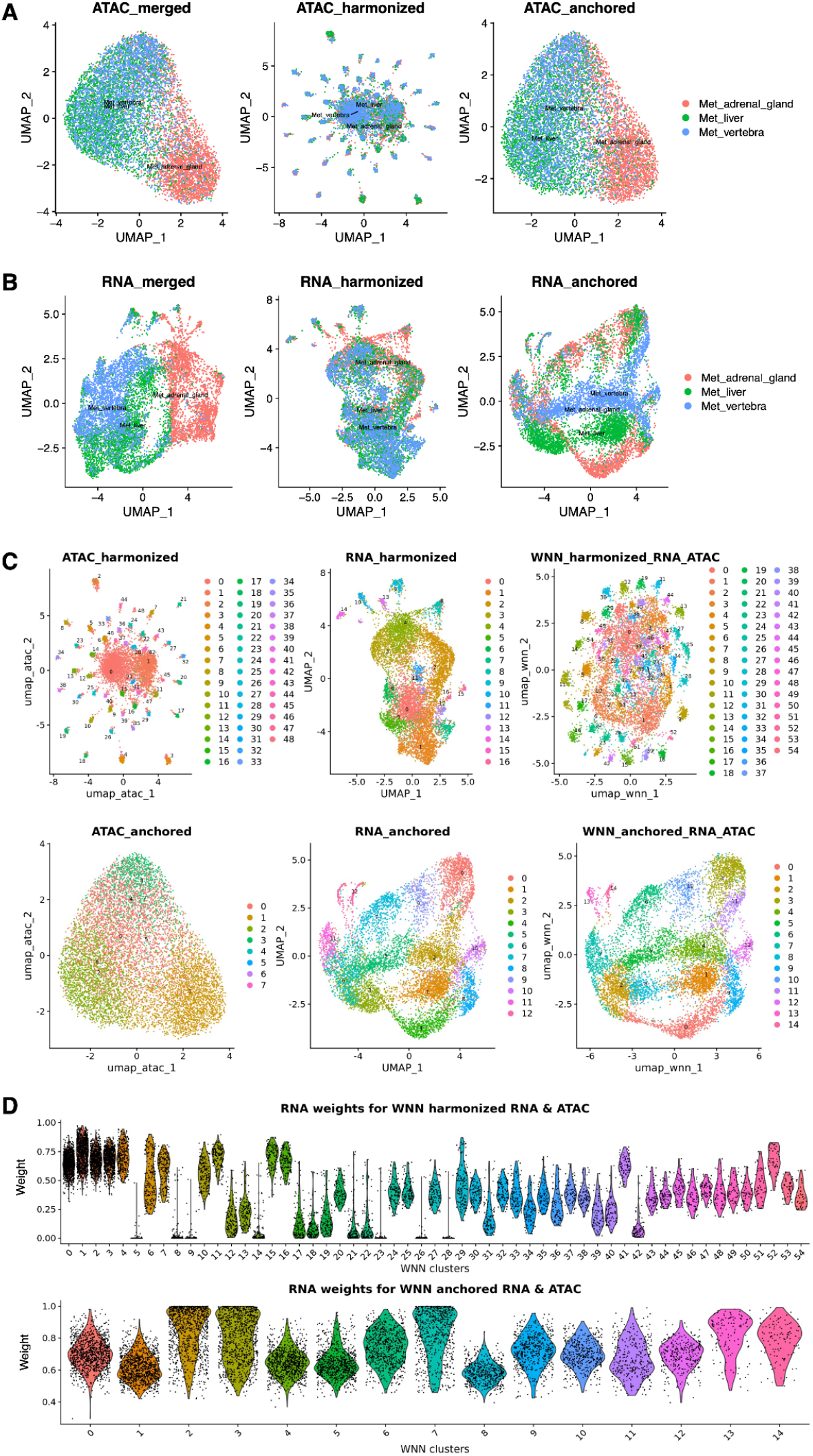
UMAP plots colored and labeled by sample IDs based on merged (left), Harmony (middle) and Seurat anchor-based integrated (right) ATAC **(A)** and RNA **(B). C)** UMAP plots colored and labeled by clusters based on RNA and ATAC data separately and integrated based on harmonized (top) and anchored (bottom) samples. **D)** Weights contributed by RNA for each WNN integrated cluster with harmonized (top) and anchored (bottom) individual scMultiome data across samples.

**Supplementary Figure 9.**
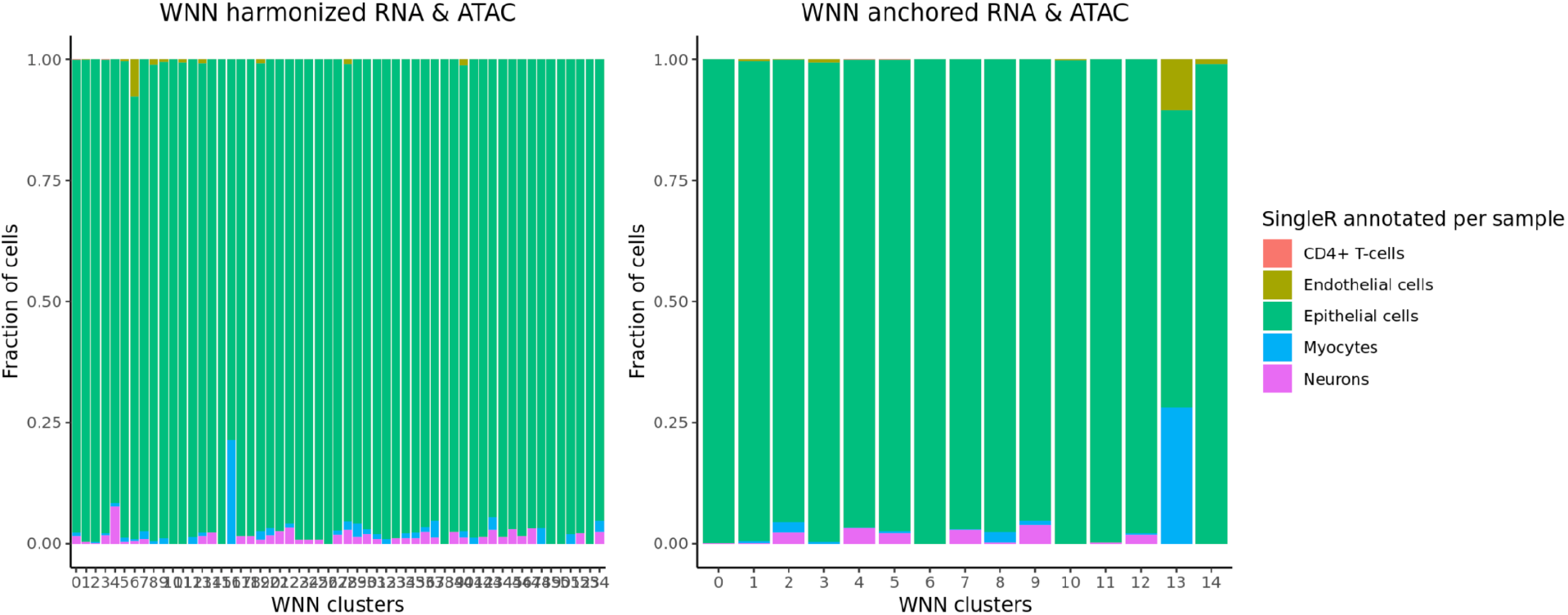
Fraction of annotated cell types across integrated samples, using Harmony (left) and Seurat anchor-based strategy (right).

### Supplementary Tables

**Supplementary Table 1:** Parameters for input configuration file.

| Parameters | Value | Note |
| --- | --- | --- |
| **PATHs** |  |  |
| pipe_dir | /path/to/iSHARC | Path to the iSHARC directory |
| work_dir | /path/to/working/dir | Desired working directory |
| **Cell Ranger ARC** |  |  |
| arc_perf | True or False | Starting with Cell Ranger ARC processed data or not. |
| arc_dir | /path/to/pre-installed/cellranger-arc | Required if arc_perf = False, and pre-installation by user is needed |
| arc_ref | /path/to/pre-downloaded/cellranger-ar/refdata | Required if arc_perf = False, and pre-download corresponding Cell Ranger ARC reference data by user is needed |
| **Samples information** |  |  |
| samples | /path/to/samples.tsv | Includes sample_id, sample_seq_id, gex_seq_path, atac_seq_path and arc_outs_path |
| integration | True or False | Whether to apply multiple samples integration |
| samples_integr | /path/to/sample_integration.tsv or NA | Includes sample_ids to be integrated |
| **Analyses** |  |  |
| second_round_filter | True or False | Whether to apply second round QC filtering |
| second_round_cutoffs | nCount_RNA_min: 1000<br>nCount_RNA_max: 25000<br>nCount_ATAC_min: 5000<br>nCount_ATAC_max: 70000<br>pct_MT_max: 20<br>TSS_Enrichment_min: 1<br>Nucleosome_Signal_max: 2 | Second round QC filtering is based on preset cutoffs and corresponding median absolute deviation. More details refer to the demo HTML reports in iSHARC/test |
| regress_cell_cycle | True or False | Whether to apply cell cycle correction |
| clustering_params | knn_k: 20<br>dims_n: 50<br>comm_res: 0.8 | key clustering parameters:<br>> knn_k: k for the k-nearest neighbor algorithm<br>> dims_n: number of dimensions (e.g., PCs) used for functions RunUMAP, FindNeighbors, FindMultiModalNeighbors.<br>> comm_res: the resolution for the function of FindClusters, value above (below) 1.0 if you want to obtain a larger (smaller) number of communities. |
| **Parallelization** |  |  |
| threads | integer (12 by default) | Number of threads for parallelization |
| future_globals_maxSize | Integer (12 by default) | The memory size in GB for global variables for the future package |

**Supplementary Table 2:**
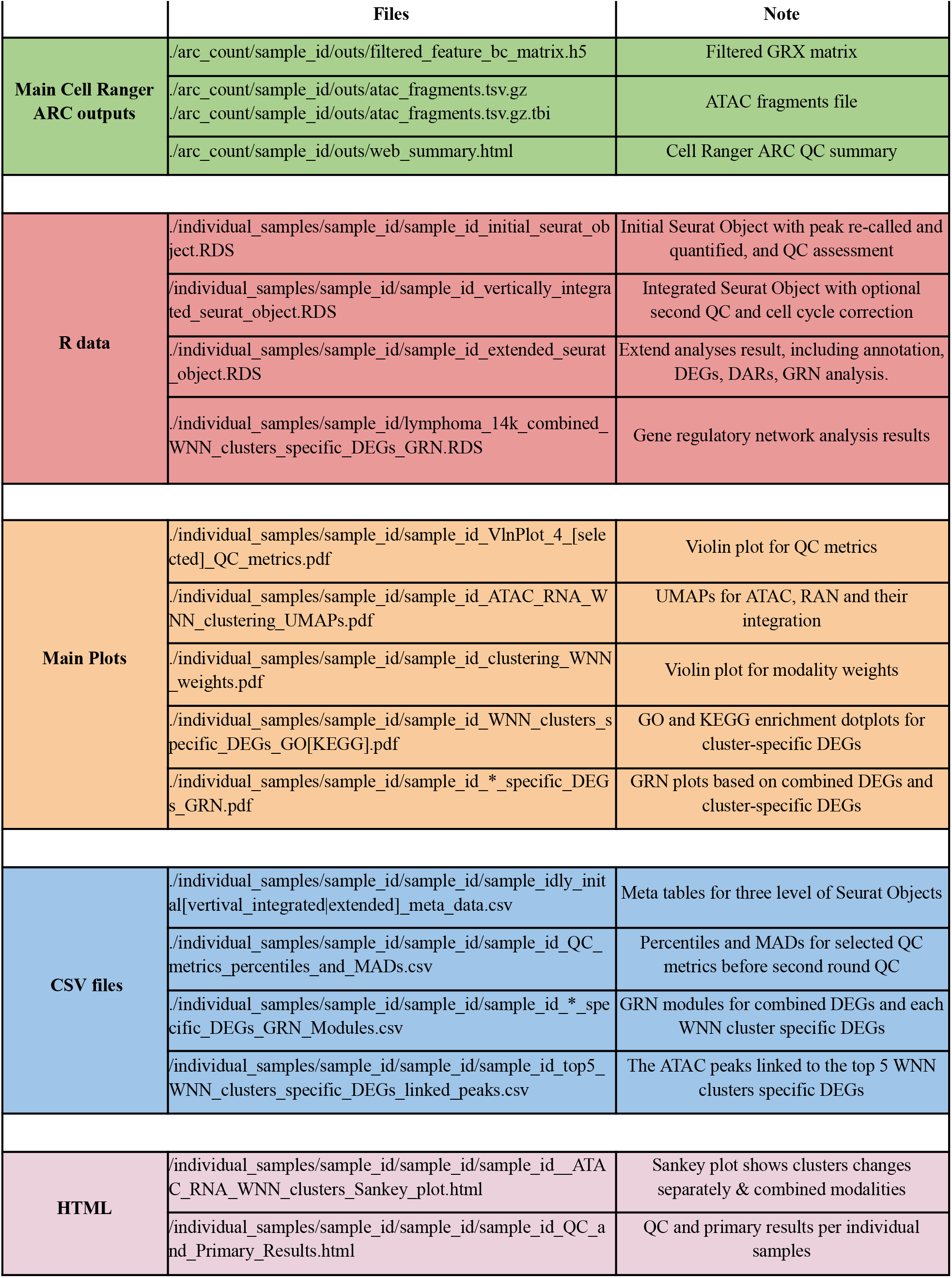
Primary output files per individual sample.

**Supplementary Table 3:**
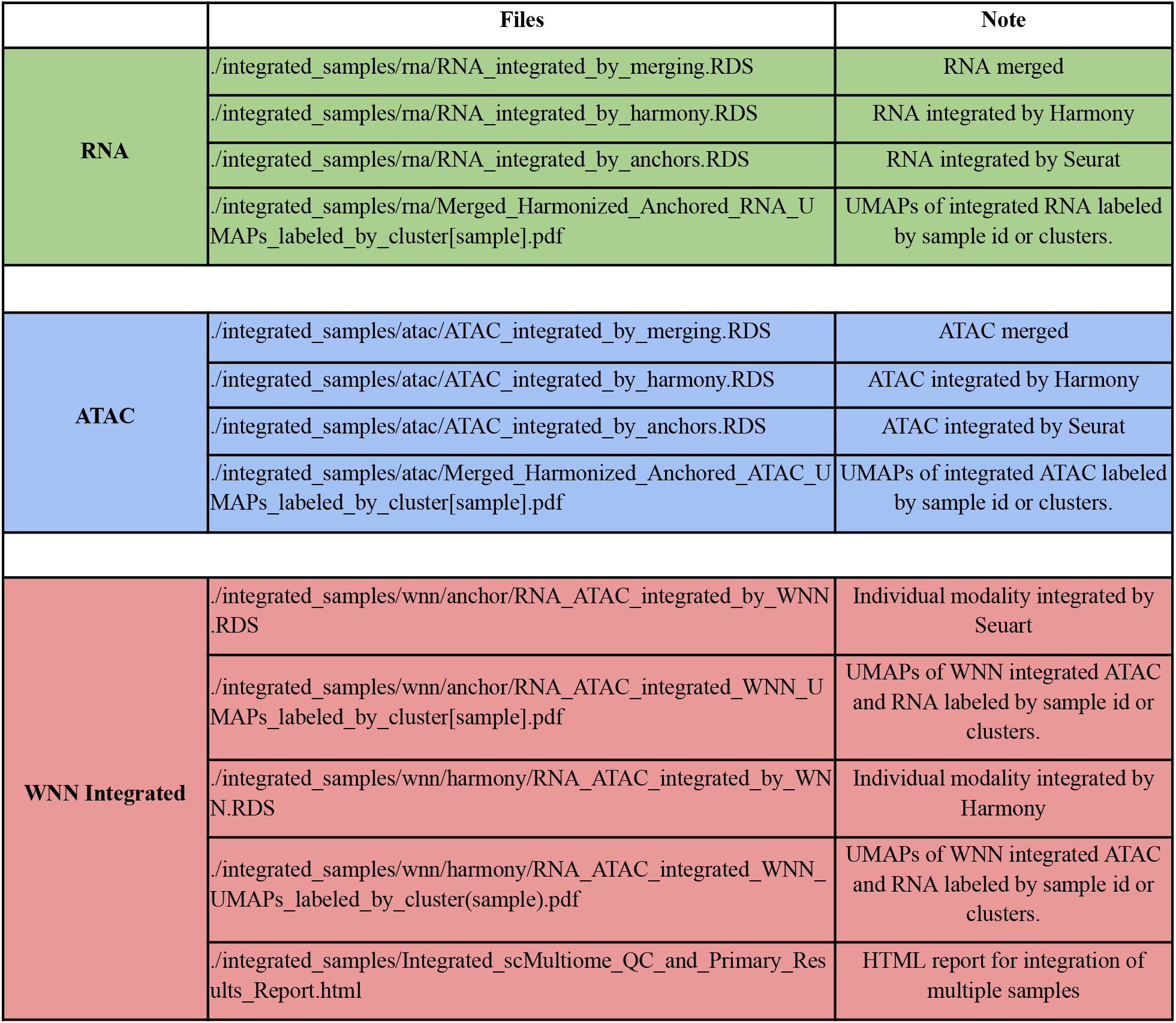
Primary output files for integrated multiple samples.

|  | Files | Note |
| --- | --- | --- |
| **RNA** | ./integrated_samples/rna/RNA_integrated_by_merging.RDS | RNA merged |
|  | ./integrated_samples/rna/RNA_integrated_by_harmony.RDS | RNA integrated by Harmony |
|  | ./integrated_samples/rna/RNA_integrated_by_anchors.RDS | RNA integrated by Seurat |
|  | ./integrated_samples/rna/Merged_Harmonized_Anchored_RNA_UMAPs_labeled_by_cluster[sample].pdf | UMAPs of integrated RNA labeled by sample id or clusters. |
| **ATAC** | ./integrated_samples/atac/ATAC_integrated_by_merging.RDS | ATAC merged |
|  | ./integrated_samples/atac/ATAC_integrated_by_harmony.RDS | ATAC integrated by Harmony |
|  | ./integrated_samples/atac/ATAC_integrated_by_anchors.RDS | ATAC integrated by Seurat |
|  | ./integrated_samples/atac/Merged_Harmonized_Anchored_ATAC_UMAPs_labeled_by_cluster[sample].pdf | UMAPs of integrated ATAC labeled by sample id or clusters. |
| **WNN Integrated** | ./integrated_samples/wnn/anchor/RNA_ATAC_integrated_by_WNN.RDS | Individual modality integrated by Seurat |
|  | ./integrated_samples/wnn/anchor/RNA_ATAC_integrated_WNN_UMAPs_labeled_by_cluster[sample].pdf | UMAPs of WNN integrated ATAC and RNA labeled by sample id or clusters. |
|  | ./integrated_samples/wnn/harmony/RNA_ATAC_integrated_by_WNN.RDS | Individual modality integrated by Harmony |
|  | ./integrated_samples/wnn/harmony/RNA_ATAC_integrated_WNN_UMAPs_labeled_by_cluster(sample).pdf | UMAPs of WNN integrated ATAC and RNA labeled by sample id or clusters. |
|  | ./integrated_samples/Integrated_scMultiome_QC_and_Primary_Results_Report.html | HTML report for integration of multiple samples |

